# Efficient exploration of peptide libraries using active learning with AlphaFold-based screening

**DOI:** 10.64898/2026.04.16.718999

**Authors:** Jokent Gaza, Jherome Brylle Woody Santos, Bhumika Singh, Ramón Alain Miranda-Quintana, Alberto Perez

## Abstract

We previously showed that AlphaFold2 can be used to screen for peptide-binding epitopes targeting the extraterminal (ET) domain of Bromodomain and Extraterminal (BET) proteins from candidate protein partners identified in pull-down experiments. However, such approaches require large numbers of AlphaFold2 calculations, making exhaustive screening impractical for larger datasets, such as viral proteomes that may target the ET domain. In many cases, identifying a substantial fraction of binders—even without exhaustive coverage—would already provide valuable biological insight into these interaction networks. Here, we show that an active learning strategy based on Thompson sampling (TS) can efficiently explore peptide sequence space. Using a library derived from BRD3 pull-down experiments, TS recovers 50% of all binders using 15% of the queries required by exhaustive sampling (3.3 times improvement over random sampling). Moreover, TS consistently identifies experimentally known binding epitopes with substantially fewer queries. Because the approach relies only on binary labels, it is readily transferable to other protein–peptide systems where AF-based binding classification is applicable, as well as to peptide-property predictors for properties such as solubility or aggregation propensity.

## Introduction

Protein–protein interactions (PPIs) orchestrate a vast number of processes within cells and organisms.^1,2^ Many of these interactions occur through short sequences in one protein– referred to as peptide binding epitopes–that recognize and bind to a specific site on another protein.^3–5^ The central role of PPIs in disease and the potential for targeting them therapeutically to interrupt pathological signaling cascades are well established.^6,7^ However, the *shallow and extended* nature of these interfaces often makes them difficult to target with small molecules.^6^ Understanding peptide–protein interfaces can therefore inform the design and optimization of small molecules or peptidomimetics.^8^ A major challenge, however, is the vast size of peptide sequence space. For example, a 12-residue peptide spans 20^12^ possible sequences. Efficient strategies that prioritize promising candidates are therefore essential for discovery.

Historically, computational screening of peptide binders has been challenging because many peptide–protein interactions involve peptides that are intrinsically disordered in solution and adopt a defined structure only upon binding. Classical docking algorithms often struggle to accurately model such folding-upon-binding processes, leading to unreliable scoring and ranking of candidate peptides. ^9^ As a result, large-scale virtual screening strategies that are common in small-molecule discovery have been more difficult to apply to peptide binders. In addition, although high-throughput experiments such as pull-down assays can suggest candidate interaction partners, they do not generally establish whether binding is direct or identify the specific peptide epitope involved.^10–12^ Recent advances in AI-based structure prediction, particularly AlphaFold2, have substantially improved the ability to model peptide–protein complexes in systems where the binding interface is sufficiently constrained.^13,14^ For example, AlphaFold competitive binding assay (AF-CBA) can be used to evaluate whether candidate peptide sequences bind to a target protein by comparing predicted structural interactions.^15,16^ These approaches make it possible to computationally screen peptide libraries for potential binders prior to experimental validation and to generate informative binary labels for adaptive search strategies.

Despite these advances, exhaustive screening remains computationally expensive. Each candidate peptide requires multiple structure predictions, and large peptide libraries may contain thousands or millions of sequences. When exploring broader sequence spaces—such as searching for viral proteins that target host domains—exhaustive enumeration becomes impractical. In many practical scenarios, identifying a substantial fraction of binders efficiently is more valuable than attempting exhaustive coverage of sequence space. Active learning strategies aim to prioritize the evaluation of candidates that are most likely to yield informative results. In particular, multi-armed bandit algorithms provide a framework for balancing exploration of uncertain regions with exploitation of promising candidates.^17,18^

Thompson sampling is a Bayesian strategy for solving multi-armed bandit problems in which actions are selected according to samples drawn from posterior reward distributions.^19,20^ This approach naturally balances exploration and exploitation and has been widely used in optimization and small molecule discovery.^21,22^ However, its application to large-scale exploration of peptide sequence space has not been explored. In this work, we formulate peptide binder discovery as a bandit exploration problem by grouping peptide sequences into clusters that represent arms with potentially different binder enrichment.

Using labels generated from our previous AF-CBA study^16^ on proteins from BRD3 pulldown experiments targeting the extraterminal (ET) domain of BET proteins,^23,24^ we apply Thompson sampling to prioritize clusters that are more likely to contain binders while still exploring uncertain regions of sequence space. The algorithm recovers 50% of all binders using 3.3 times fewer queries than random sampling and consistently identifies known binding epitopes earlier during the search. Because the approach relies only on binary binding labels generated from structure prediction assays, it can be readily applied to other protein–peptide systems where AlphaFold-based screening is feasible or to peptide-property predictors such as solubility and aggregation propensity. Efficient exploration strategies such as the one presented here will become increasingly important as structure prediction enables screening of ever larger peptide and protein libraries.

## Methods

### Motivation

Thompson sampling is a strategy for solving multi-armed bandit problems,^25^ often illustrated using casino slot machines. Each arm (or slot) has an unknown probability of payout, and the objective is to maximize cumulative reward over repeated plays by balancing exploration (testing uncertain arms) and exploitation (favoring promising arms). Instead of assigning fixed estimates for the reward, TS maintains a probability distribution over the expected reward of each arm. For binary outcomes (winning coins vs losing coins), this uncertainty is naturally modeled using Beta distributions where each arm is parameterized by *α* (number of wins) and *β* (number of losses). These distributions, thus, represent the posterior belief over the success probability of each arm. In the TS workflow, an agent samples a plausible reward rate 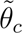 from every arm’s Beta distribution and selects the arms with the highest sampled values. After selecting slots to play and observing the outcomes, the agent performs a Bayesian update by adjusting the *α* and *β* parameters of the corresponding distributions. Over time, this sampling-based decision rule automatically concentrates selections on high-reward arms while continuing to explore uncertain ones.

### Thompson Sampling Workflow

We aim to use the above strategy to identify peptide likely to bind a particular protein – from a pool of candidates that would be too computationally demanding to explore exhaustively. Here, sequence-clusters will be analogous to the casino slots where, instead of winning or losing coins, we obtain binder or non-binder sequences upon querying. Using the results from the previous AF-CBA exhaustive study,^16^ we constructed a dictionary of binder sequences from human proteins, which were then clustered (see next sections). Each sequence is labeled in binary (binder = 1, non-binder = 0). Formulated as a multi-armed bandit problem, the objective is to maximize the number of binders discovered under a limited number of queries (i.e., AF2 runs or, in this case, a dictionary lookup) by using Thompson sampling to decide which clusters to query based on their inferred Beta distributions.

We start the workflow by constructing the initial priors for each cluster *c* (Figure 1). Our hypothesis is that clusters differ in their binder prevalence, and that some clusters contain a higher fraction of binder sequences (relative to their total size) than others. In particular, binders are expected to be concentrated within specific clusters, which should therefore exhibit posterior distributions shifted toward higher success probabilities. The prevalence of binder sequences within cluster *c* is represented by the success probability *θ_c_* of a Bernoulli outcome, with prior *θ_c_* ∼ Beta(*α*_0_*, β*_0_). We found that initializing *α*_0_ and *β*_0_ based on the global hit rate of the full dictionary (3,393 / 142,338 ∼ 2.4%) led to a more efficient TS (Figure S2). For new proteomes, this rate is unknown. However, since non-binders are expected to dominate, initializing with *β*_0_ *> α*_0_ seems to be a reasonable default.

**Figure 1:**
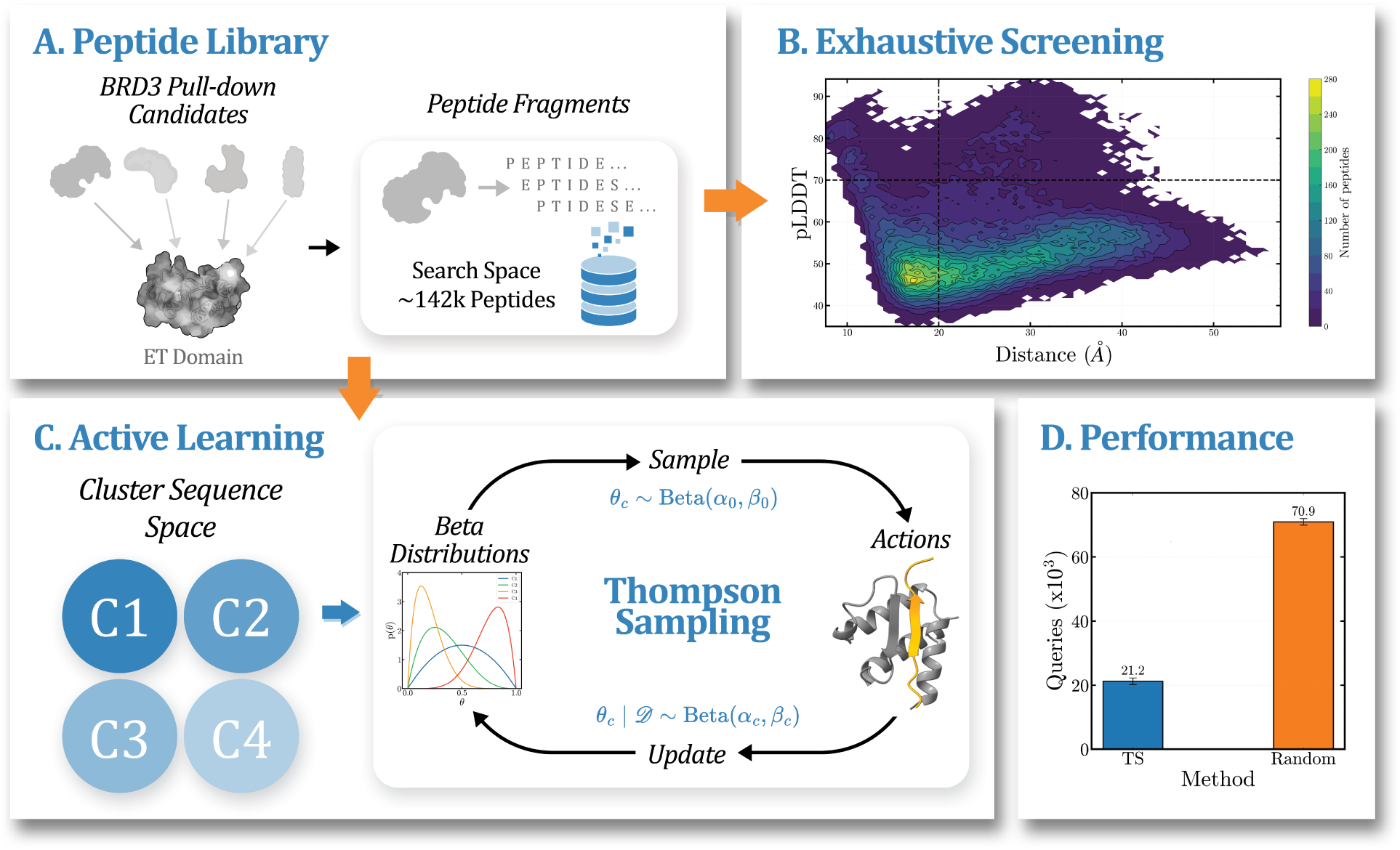
(A) Candidate proteins from pull-down experiments are fragmented into 25-AA peptides using a 1-AA sliding window for a total of 142,338 peptides. (B) Exhaustive screening for binders is performed by runnning one AlphaFold2 prediction for each ET:peptide complex. (C) Active learning approach using Thompson sampling where each cluster is assigned a Beta distribution. For each iteration, these distributions are sampled and clusters with the highest values are selected. Labels for a set number of peptides from the selected clusters are then revealed and used to update the posterior Beta distributions. (D) To recover 50% of the binders, TS requires sampling only 15% of the total sequences, a 3.3-times improvement over random sampling.

To reduce sensitivity to the chosen priors and to refine these prior distributions, we introduce a random *seed* set consisting of sequences sampled uniformly from the full library. All seed labels are revealed and used to update the Beta distributions (*θ_c_* | 𝒟 ∼ Beta(*α_c_, β_c_*)) for clusters represented in the seed: *α_c_* = *α*_0_ + *p_c_, β_c_* = *β*_0_ + *n_c_*, where *p_c_* and *n_c_* denote the numbers of binders and non-binders observed in cluster *c*, respectively. The same posterior update rule is applied after each subsequent selection round in the actual TS rounds.

Active selection then proceeds in rounds with a fixed batch size (which we set to 50 as a balance between computational expense and getting sufficient knowledge to update distributions). At each round, Thompson sampling is applied by drawing one sample from each cluster posterior, 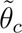 ∼ Beta(*α_c_, β_c_*), which represents a plausible binder rate under current knowledge. Because cluster sizes are highly heterogeneous (including singleton clusters), clusters with no remaining unqueried sequences are excluded from consideration in that round.

At each round, clusters are ranked by their Thompson samples 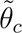 and the top *k* non-exhausted clusters are selected for allocation. The batch budget is then distributed across these clusters using either *equal* or *proportional* allocation. Under equal allocation, each selected cluster receives an identical query quota. Under proportional allocation, cluster quotas are assigned in proportion to 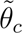, favoring clusters with higher sampled binder prevalence. For each selected cluster, peptides are chosen randomly from the set of unqueried peptides up to the allocated quota for that cluster. When a cluster cannot satisfy its full quota (due to limited remaining peptides), the unused budget (i.e. the *leftovers*) is redistributed to lower-ranked clusters in the global 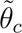 ordering that still contain unqueried peptides. If capacity remains after this redistribution, the batch is topped up by iteratively sampling single peptides from the best available clusters until either the 50 selections are made or all clusters are exhausted. Labels for all selected peptides are then revealed and for each cluster touched in the round, the posterior parameters are updated using the method described previously.

We evaluate the performance of TS using a discovery curve that tracks the cumulative number of binders as a function of cumulative queries. These results are compared to a baseline run with the same initialization seed *S*, batch size, and RNG seed, but using uniform random sampling from the full dictionary. This provides a matched control for quantifying the benefit of cluster-aware Thompson sampling. For reproducibility, all stochastic components (seed initialization, posterior sampling, within-cluster sampling, and refill) are driven by a single pseudorandom generator initialized from a user-specified seed, enabling exact replay of selection trajectories.

### Building the Dictionary

To construct the peptide epitope dataset, a list of candidate BRD3-associated proteins was compiled from genes reported to co-precipitate with FLAG-tagged BRD3 in pull-down and biophysical assays, yielding 335 candidates.^26^ Corresponding human protein sequences were retrieved from UniProt^27,28^ and restricted to reviewed (Swiss-Prot) entries. After removal of unavailable or redundant records, the final dataset comprised 318 proteins (see SI), including gene names, UniProt IDs, and full-length sequences.

The BRD3 ET domain is a three-helical bundle with a flexible loop that undergoes a disorder-to-order transition upon peptide binding.^29^ Structural studies indicate that a peptide length of 25 amino acids favors stable binding within the ET binding pocket. Accordingly, we generated a peptide fragment library from each full-length protein sequence using a 1-AA sliding window approach. Overlapping 25–amino acid peptides were produced with a single-residue step size across the entire sequence, thereby generating all possible 25-mer fragments for each protein.

Each peptide fragment was subsequently modeled in complex with the BRD3-ET domain using ColabFold using their sequence as input.^13,30^ For every BRD3–peptide pair, five structural models were generated. To classify peptides as binders or non-binders, we applied a two-parameter filtering strategy. First, structural confidence was assessed using the mean peptide pLDDT score, requiring a value greater than 70. Second, spatial proximity was evaluated by calculating the average *C_α_* − *C_α_* distance between peptide residues and key binding residues (I42, E43, and I44) of the ET domain, with a threshold of less than 20 °A. To better capture confidence within the likely binding region, rather than averaging pLDDT and distance over the entire peptide in all cases, we estimated these values within a putative binding segment when possible. Because the binding region is unknown *a priori*, we identified consecutive residue runs with pLDDT ≥ 70 and length ≥ 6 residues. If such a segment existed, the mean pLDDT and distance over this region were used. Otherwise, the values over the full peptide were used. This procedure accounts for binders with long non-interacting tails.

All five predicted models were evaluated for each peptide, and a peptide was classified as a binder if at least four out of five models (≥ 80%) satisfied both criteria. Peptides failing to meet this threshold were classified as non-binders. Finally, results from all proteins were aggregated, and duplicate peptide sequences were removed to generate a non-redundant peptide dictionary. The resulting dataset comprises 142,338 unique 25-mer peptide sequences derived from proteins associated with BRD3 ET complex formation, forming the basis for subsequent clustering and TS analyses.

### Clustering

To partition the full peptide dictionary into groups of related sequences, we clustered the full set of unique 25-mer peptides using three sequence-based procedures: CD-HIT (v4.8.1),^31^ the MMseqs2 easy-linclust, and MMseqs2 easy-cluster workflows (v17.b804f).^32^ All methods were run at sequence identity thresholds of 0.4, 0.5, 0.7, and 0.9. These approaches were selected to compare a widely used sequence identity-based clustering approach (CD-HIT) with two MMseqs2 workflows that differ in sensitivity and computational cost. Throughout this work, clustering generated by MMseqs2 easy-linclust is referred to as LINCLUST, whereas clustering generated by MMseqs2 easy-cluster is referred to as LINCLUST/MMseqs2.

CD-HIT clustering was performed with word sizes chosen according to the recommended parameter ranges for each threshold. In this setting, the CD-HIT identity thresholds define the minimum sequence similarity required for assignment to a cluster, and each cluster is represented by a single representative sequence.

For MMseqs2-based clustering, the minimum sequence identity and minimum coverage threshold were set to the same value at each clustering level, and the default coverage mode was used throughout. Thus, as the threshold increased, both sequence identity and alignment coverage became stricter for each MMseqs2 run.

The LINCLUST workflow corresponds to the standalone linear-time clustering procedure implemented in MMseqs2 and was used to obtain a fast partitioning of the peptide library based on sequence similarity. In contrast, LINCLUST/MMseqs2 refers a combined cascaded clustering workflow that extends Linclust with the more sensitive MMseqs2 clustering procedure, thereby increasing sensitivity relative to LINCLUST alone at the expense of additional computation.

## Results and discussion

### Performance against Random Sampling

Since each peptide query requires an AlphaFold2 structure prediction, the practical goal of Thompson sampling is not exhaustive enumeration of sequence space, but maximizing binder discovery under a limited query budget. We therefore first optimized the TS hyperparameters controlling initialization seed size *S*, the number of selected clusters *k*, and the Beta prior parameters *α*_0_ and *β*_0_ (Figures S2-S4). Results are presented as discovery curves for the full dataset, tracking the number of binders discovered versus the total number of peptides queried. A more efficient algorithm discovers more binders in fewer rounds and should outperform uniform random sampling.

We found that larger values of *S* and *k* generally produced steeper TS discovery curves. Early-round performance also depends on the choice of prior. TS runs initialized with a uniform Beta prior for all clusters showed poorer performance. This occurs because such priors change slowly after only a few observations, leaving clusters without binders nearly as likely to be selected as untested clusters. Regarding allocation strategy, we observed that *equal* and *proportional* allocation produced similar results (Figures S3,S4). For the remainder of the study, we therefore report results using *proportional* allocation.

From the hyperparameter screening, we assumed that the effects of each hyperparameter were approximately additive. For a batch size of 50, the optimized parameters were *S* = 5000*, k* = 50, and prior parameters *α*_0_ = 0.2 and *β*_0_ = 9.8. We then ran 100 TS replicates using these parameters on clusters generated by CD-HIT, LINCLUST/MMseqs2, and LINCLUST (Figure 2). For all runs across clustering methods, we quantified performance using the area under the discovery curve (AUC). We observed similar results for clustering parameters 0.4, 0.5, and 0.7, but worse performance at 0.9. This likely reflects differences in cluster granularity, as higher sequence identity values produce more clusters. This trend is expected because TS performs best when the dataset is well discriminated, with most binders concentrated in a relatively small subset of clusters. Because performance was comparable for 0.4, 0.5, and 0.7, we focus on the results obtained with parameter 0.5 for the remainder of the paper.

**Figure 2:**
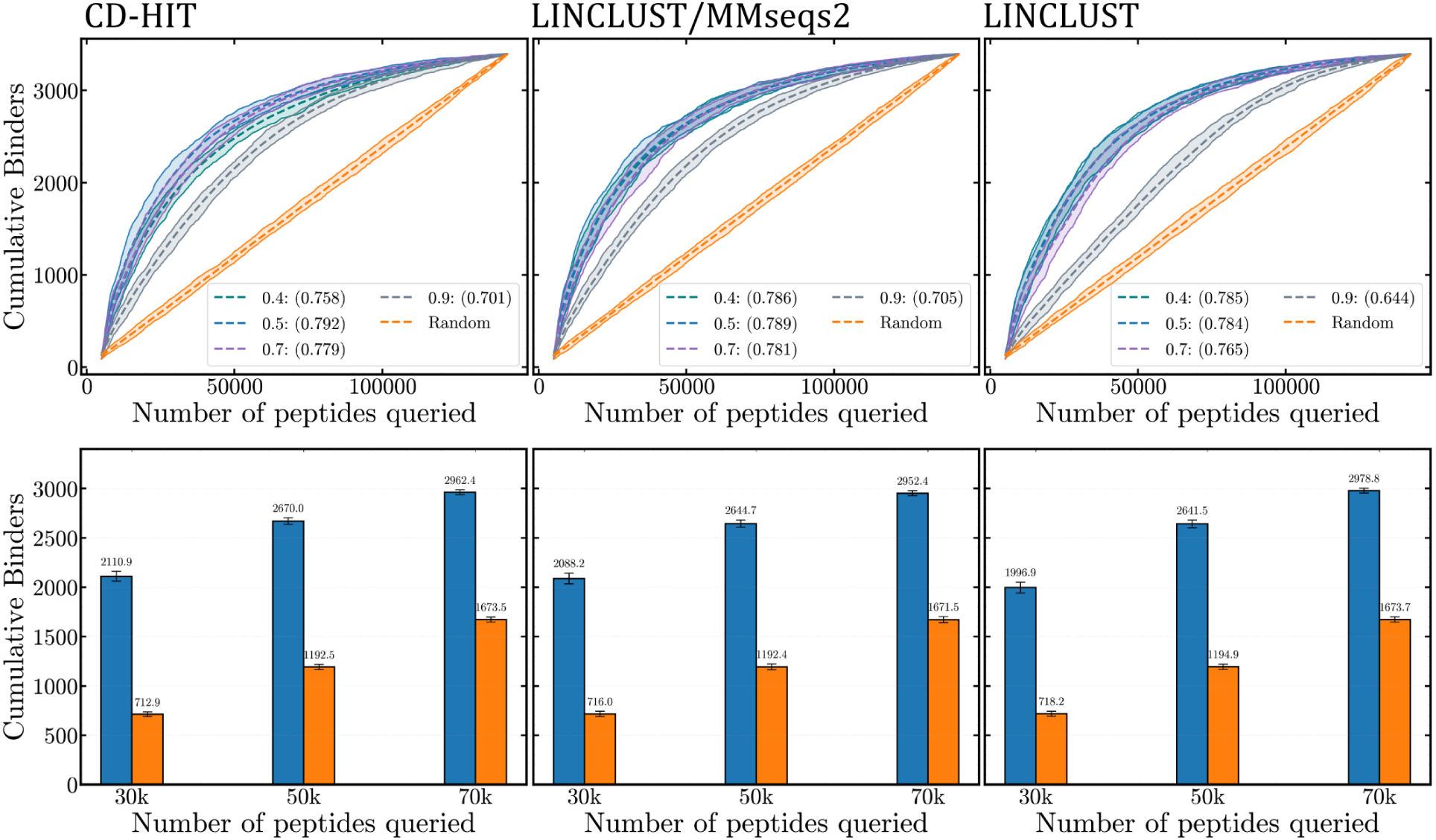
Performance Summary of Thompson Sampling vs. Random Sampling. (Top) Cumulative binders sampled relative to the total number of peptides checked across all clustering algorithms (CD-HIT, LINCLUST/MMseqs2, and LINCLUST). Results are aggregated over 100 replicates. For each clustering method, the mean (dashed line) and the min/max results (solid lines) define the shaded region, representing the worst- and best-case scenarios. The normalized area under the curve (AUC) is reported (in parenthesis) to quantify TS performance. (Bottom) Cumulative binders identified for clustering algorithms using a 0.5 parameter at checkpoints of 30k, 50k, and 70k total peptides checked.

To quantify the practical value of active learning, we next evaluated binder recovery at fixed query budgets of 30k, 50k, and 70k peptides, corresponding to approximately 20%, 35%, and 50% of the full dataset, respectively. At these checkpoints, TS was approximately 2.9-, 2.2-, and 1.78-times more efficient than random sampling, respectively. These budgets recovered approximately 62%, 78%, and 87% of the 3393 binders in the dataset.

### Clustering Results

To understand why TS performs best for some clustering settings, we next examined how the peptide library is partitioned into clusters, which serve as the arms in the bandit formulation. We first visualized the overall distribution of binders and non-binders by plotting the pLDDT against the average distance to the BRD3 binding site (Figure 3). Binder classification is based on the five AF2 models generated per peptide: each model yields a pLDDT–distance pair, and a peptide is labeled as a binder if at least four of the five pairs satisfy the thresholds (pLDDT ≥ 70 and distance *<* 20 °A). Although many predictions exhibit high pLDDT values at large distances from the active site, these do not generally correspond to alternative binding modes. Instead, they typically represent peptides that adopt some degree of secondary structure, as reflected by their low radius of gyration, while remaining spatially separated from the ET receptor.

**Figure 3:**
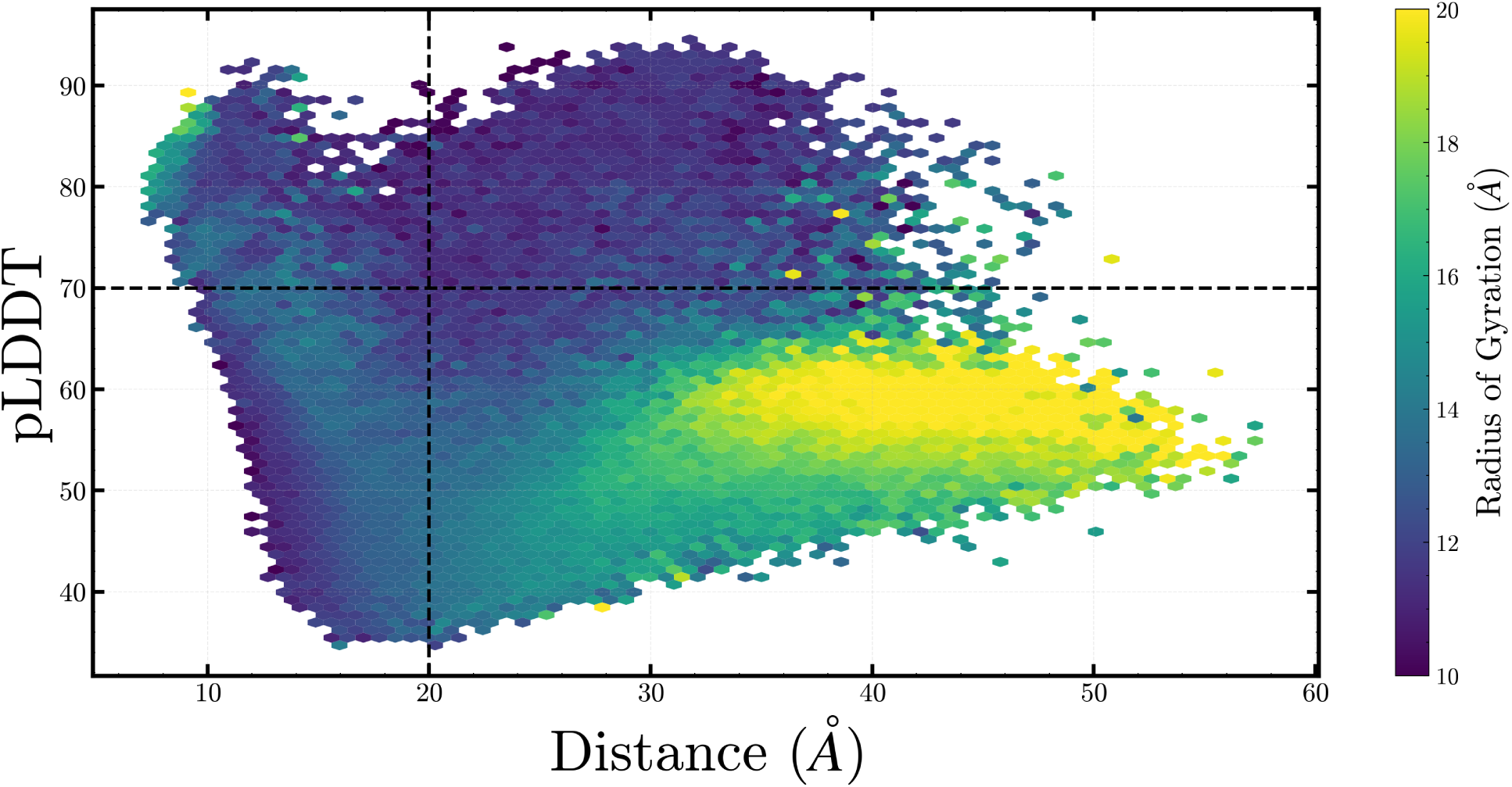
AF2 predictions of each peptide in complex with the BRD3 target. Each AF2 run produces five models, which are evaluated by pLDDT and distance to the binding site. A sequence is classified as a binder if at least four of the five models have pLDDT ≥ 70 and distance *<* 20 °A. The distribution is visualized as hexagonal bins, aggregating peptides by mean pLDDT and distance, with color mapping representing the mean radius of gyration of the five models.

This distribution mainly serves to visualize how binders and non-binders can be separated. By clustering peptide sequences (see methods^31,32^), we transform the exploration of sequence space into a bandit problem where each cluster represents a hypothesis about binder enrichment. For each clustering algorithm, we generated four different cluster sets using different sequence identity cutoffs. This user-defined identify levels controls how similar sequences must be in order to be grouped into the same cluster. We found that increasing this parameter produces a larger number of smaller, sparse clusters (Figures 4A). In contrast, lower values produce fewer clusters, with binders concentrated in a small subset of them. For example, using a threshold of 0.5 (Figure 4B), we found that, on average, 50% of all binders were contained in only 151 clusters, whereas for 0.9 the same fraction required 440 clusters. This distinction helps explain the TS results above. When binders are concentrated into fewer clusters, early successes provide stronger evidence of enrichment and can be more effectively exploited in subsequent rounds. In contrast, finer-grained clustering spreads binders across many sparse clusters, thus making it harder for TS to exploit enriched regions.

**Figure 4:**
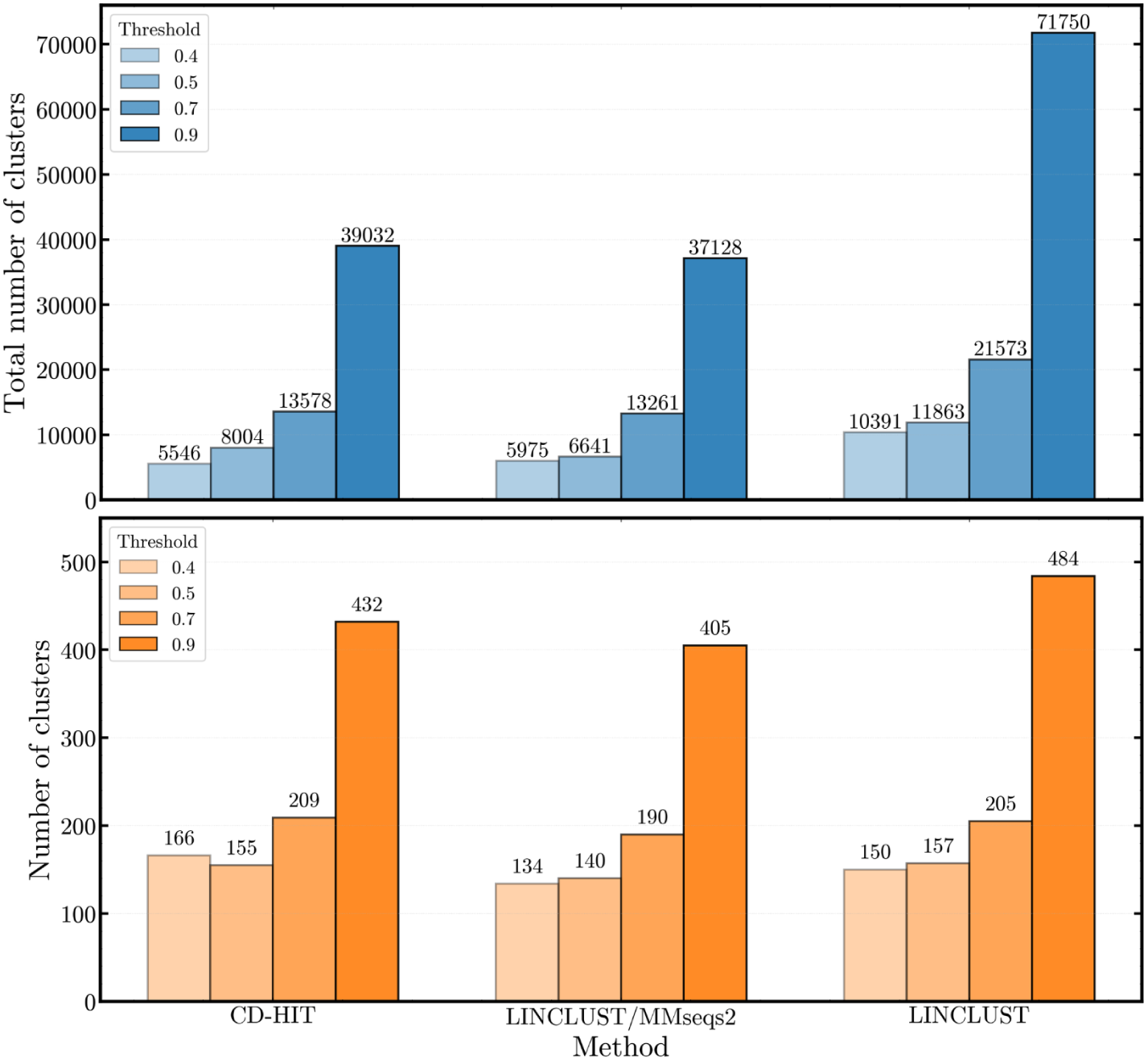
(Top) Total number of clusters generated by each method. (Bottom) Minimum number of clusters required to exhaust to capture 50% of all binders.

### Top Peptides

Although TS discovers more binders than random sampling, it is clear that not *all* binders can be identified with substantially fewer queries. The discovery curves plateau after approximately half of the dataset has been queried. We hypothesize that this behavior arises from clusters that contain binders only sparsely relative to non-binders. In such clusters, binder discovery becomes unlikely because (1) their Beta posteriors are dominated by non-binder observations, and (2) even when such a cluster is selected, identifying a binder still depends on random within-cluster sampling.

To support this explanation, we examined the unsampled binders after 70k queries (Figure S5) of one TS run. In this run, 259 out of 8004 clusters remained unexhausted after 70k queries. As expected, the remaining binders were distributed across these clusters, most of which contained fewer than 10 binders. Given the high prevalence of non-binders in these clusters, sampling from them is more likely to yield non-binders, which further shifts their Beta posteriors downward and reduces their probability of being selected in subsequent rounds.

An important question is therefore the quality of the binders identified by TS. Because of how the dictionary was constructed, many clusters contain highly similar sequences, as these peptides were generated by sliding a 1-AA window across full protein sequences. For ET, peptides interact through a conserved motif present in both human and viral proteins. Consequently, sequences containing this motif are also expected to be binders. This is particularly true for experimentally validated ET binders, which adopt an alternating hydrophilic–hydrophobic motif at the interaction site. Such known binders should therefore reside in clusters that also contain other motif-bearing sequences and enriched binder population.

To assess binder quality, we examined how many queries were required to recover five known ET binders: BRG1, INO80B, CHD4, NSD3, and BICRA. Specifically, we tracked the sequence that appeared as top candidates in the completed AF-CBA pipeline of these proteins. These epitopes bind ET at the *β*-hairpin site using one or two *β*-strands (Figure 5A).

**Figure 5:**
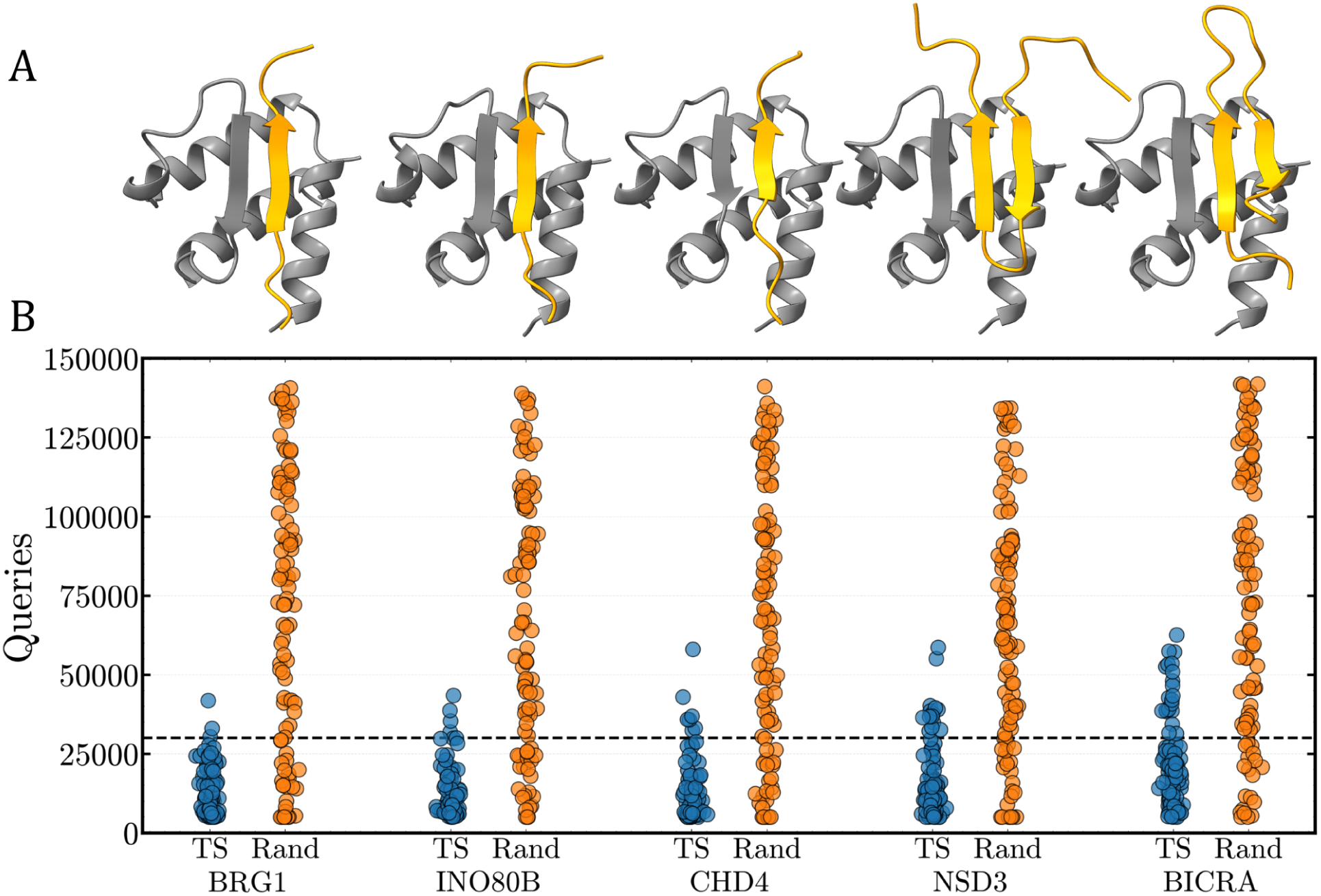
Performance of TS in identifying top peptides using the CD-HIT clustering method (sequence identity parameter 0.5). (A) AF2 predictions of the top peptides (orange) in complex with the BRD3-ET domain (gray). Unstructured regions not interacting with the ET domain are hidden for clarity. (B) Number of queries required to identify the binders BRG1, INO80B, CHD4, NSD3, and BICRA in 100 runs of TS (blue) and random sampling (orange). A visual threshold at 30k checks is indicated by the dashed horizontal line.

Across 100 TS runs using CD-HIT clustering (wih sequence identity of 0.5), BRG1, INO80B, and CHD4 were recovered with high probability after 30k queries (∼20% of the dataset; Figure 5B, S7). BRG1, INO80B, and CHD4 were identified in 97/100, 95/100 and 93/100 runs, respectively. Recovery rates for the remaining peptides were lower: 85 for CHD4, and 79 for BICRA. These differences are consistent with the cluster composition of each peptide (Figure S6). BRG1, INO80B and CHD4 reside in clusters enriched in binders, whereas the other peptides occur in clusters containing more non-binders, reducing their likelihood of selection during cluster sampling. These results show that TS does not simply improve overall binder counts, but also prioritizes biologically-known binders in early TS rounds.

### Inner workings of TS

To gain more insights into why TS is efficient, we examined the Beta distributions of two representative clusters: cluster 1 (the most populated cluster) and cluster 1105, which contains the top peptide sequence of INO80B. In terms of size, cluster 1 is much larger (177 sequences) but contains no binders, whereas cluster 1105 is smaller (25 sequences) but has a high binder fraction (19 binders).

We tracked the Beta distributions (Figure 6A,B) of these clusters as they were sampled during a TS run (CD-HIT clustering, sequence identity parameter 0.5). As initialized, both clusters start with priors skewed toward low success probability, resulting in similar early selection likelihood. However, for cluster 1, repeated sampling yields only non-binders, causing its Beta distribution to shift further left as *β_c_* increases. Consequently, its probability of selection decreases over time. In contrast, cluster 1105 rapidly accumulates binder observations, shifting its Beta distribution to the right and producing larger sampled *θ_c_* values, which increases its selection probability. This behavior explains why the INO80B sequence is typically recovered within 30k queries.

**Figure 6:**
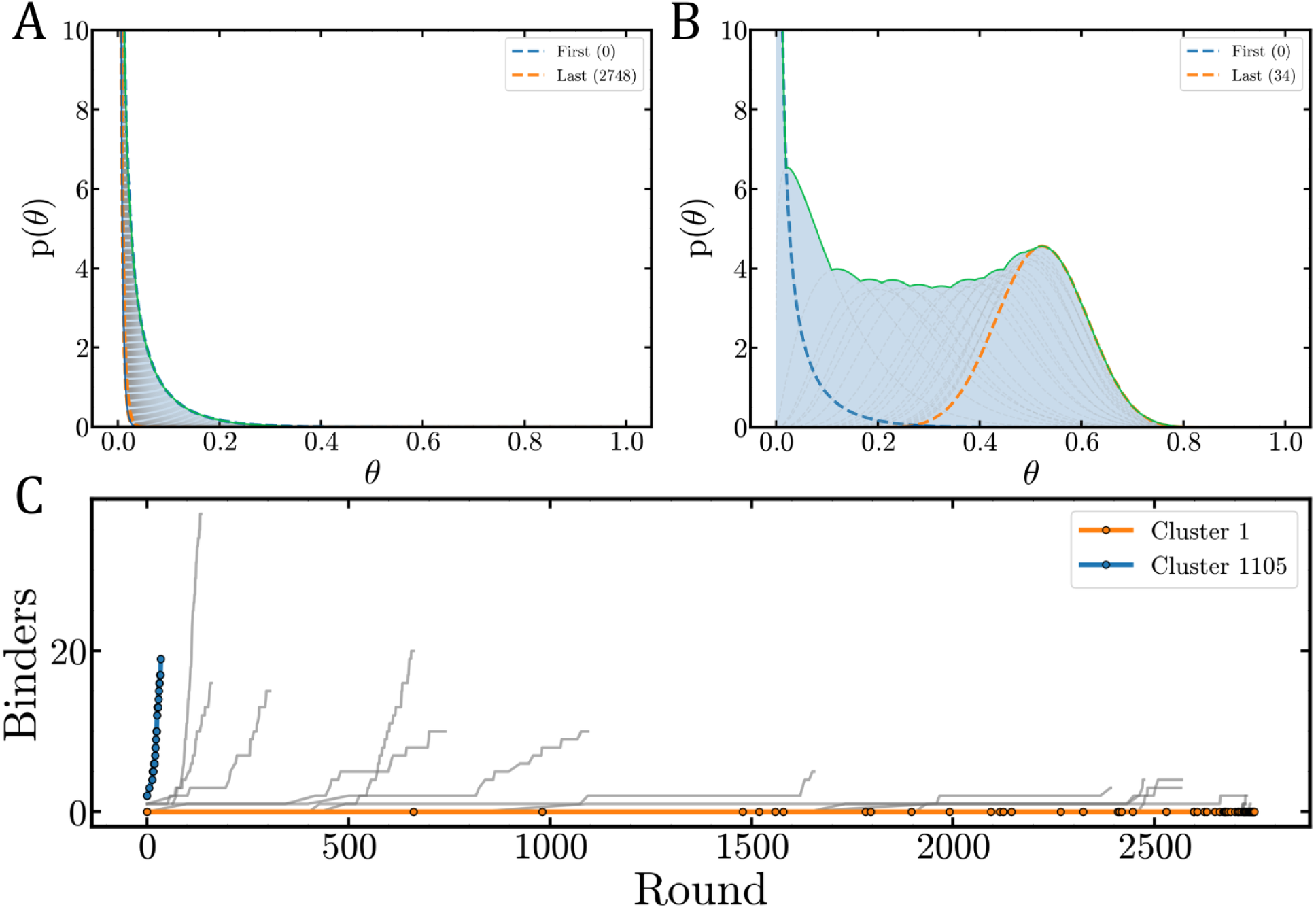
Enhanced Sampling of Clusters. Beta distributions of the (A) most populated cluster (0 binders in 177 sequences) and (B) the cluster containing INO80B (19 binders in 25 sequences) as they are updated during the TS workflow. The first and last rounds these clusters were checked are in blue and orange, respectively. (C) Sampling history of the 50 most populated clusters compared to the cluster containing INO80B (Cluster 1105).

To further illustrate TS efficiency, we examined the sampling histories of these clusters alongside the 50 most populated clusters (Figure 6C) from the same TS run. Cluster 1 is sampled across the entire run, but because its posterior shifts leftward, it is selected increasingly rarely until the later rounds. In contrast, once binders are observed in cluster 1105, its sampling frequency rises sharply until the cluster is exhausted. This pattern is also observed for other clusters with high binder population. The steep sampling trajectories of these clusters demonstrate how TS prioritizes clusters with evidence of binding activity, leading to efficient binder discovery.

Here we have exemplified the approach on the ability to bind ET, in a similar fashion to what has been done for small molecules. The approach is readily transferable to other problems were binary labeling is available, such as other protein-peptide systems where the AF-CBA can be applied. As a proof of concept, we also applied our TS strategy to the current library of 142,338 peptide candidates by labeling according to soluble or non-soluble peptides (see SI) as calculated using NetSolP,^33,34^ showing similar performance for what has been shown for binders.

## Conclusion

Recent advances in machine learning have greatly expanded the molecular search space accessible to computational chemistry. Community-driven development and the open-source nature of these models, such as AF2, have further enabled researchers to extend these tools beyond their original design and build new pipelines that exploit their strengths. We have previously shown that AF2, although not originally developed for this purpose, can be adapted to distinguish binder from non-binder peptides. Building on this idea, we also developed a pipeline to identify peptide binding epitopes within known binding proteins for target proteins of interest. Here, we asked whether this search for binding epitopes could be made more efficient using an active learning strategy.

Using Thompson sampling, we show that binders can be recovered more efficiently than by random sampling. As expected, TS performs best when applied to clusterings in which binders are concentrated within a relatively small number of clusters. In addition to improving binder recovery, TS also identifies the top epitopes more rapidly. We further find that this framework is not limited to binder discovery. When applied to NetSolP-based labels, TS similarly enriches for highly soluble peptides. Overall, these results demonstrate that TS provides an efficient strategy for prioritizing peptide sampling in sequence space and can help scale searches for binding epitopes in larger protein libraries or entire proteomes. The approach is also general and applicable to any peptide-property predictor that provides binary labels, including binding, solubility, or aggregation propensity.

## Supporting Information Available

### Data and Software Availability

The TS implementation is available in our Github repository: https://github.com/PDNALab/Thompso

### Supporting Information

TS performance on solubility data. Hyperparameter screening for seed size *S*, number of clusters *k*, and priors *α*_0_*, β*_0_. Complete results for all TS runs. Distribution of binders and non-binders in selected clusters. List of genes and their corresponding sequence.

### Authors’ contributions

J.G. performed the TS runs. J.S. built the clusters. B.S. built the protein library. R.M.Q. and A.P. conceived the project. All authors wrote, revised, and approved the final manuscript.

## Acknowledgement

A.P., J.G., and B.S. thank the National Institute of General Medical Sciences of the National Institutes of Health under award number R01GM149646. R.M.Q. and J.S. also thank the same institute under award number R35GM150620.

## TOC Graphic

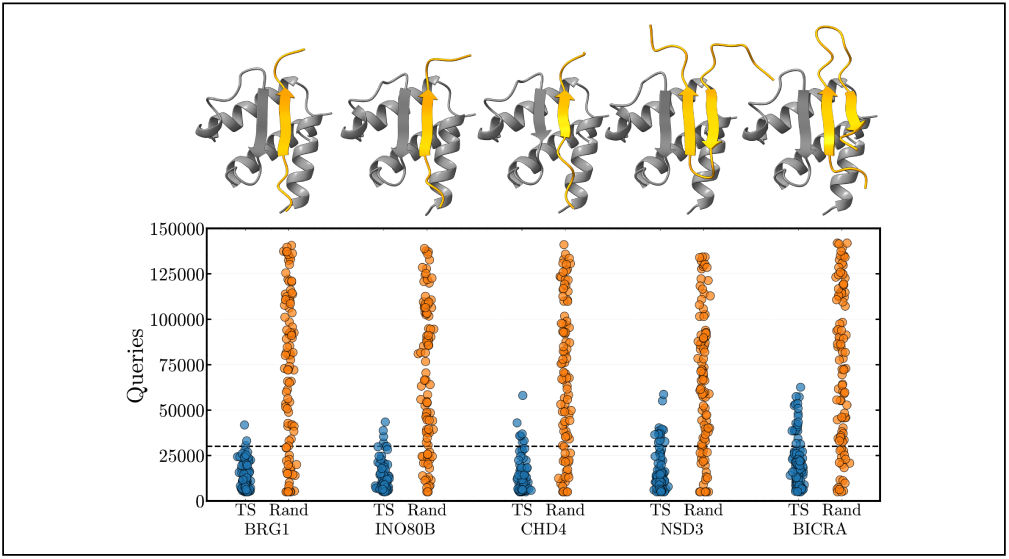

