## Supplementary Information for "Efficient exploration of peptide libraries using active learning with AlphaFold-based screening"

### TS for finding the most soluble peptides

The utility of TS is not limited to AF2-derived binder labels. Since the peptide library consists of fragments obtained using a 1-AA sliding window, many sequences share similar physicochemical properties. As an example, we asked whether TS can also efficiently identify peptides with high predicted solubility within the same clustered sequence space.

To construct a solubility-based dataset, we predicted peptide solubility using a local installation of NetSolP 1.0 (ESM-1b distilled language model).[1,2] NetSolP outputs a continuous score between 0 and 1, where values closer to 1 indicate higher predicted solubility. To enable direct comparison with the binary binder task, we converted these scores into three binary labels by defining soluble peptides as the top 2%, 5%, or 10% of sequences by predicted solubility. The peptide library was clustered using CD-HIT, and TS was applied under the same replay framework used for binder discovery (Figure S1). TS exhibited discovery curves closely matching those observed in the binder dataset, with similarly steep early enrichment and comparable normalized area under the curve. These results demonstrate that TS generalizes beyond binder identification and can efficiently prioritize peptides enriched for other sequence-derived properties, such as solubility.

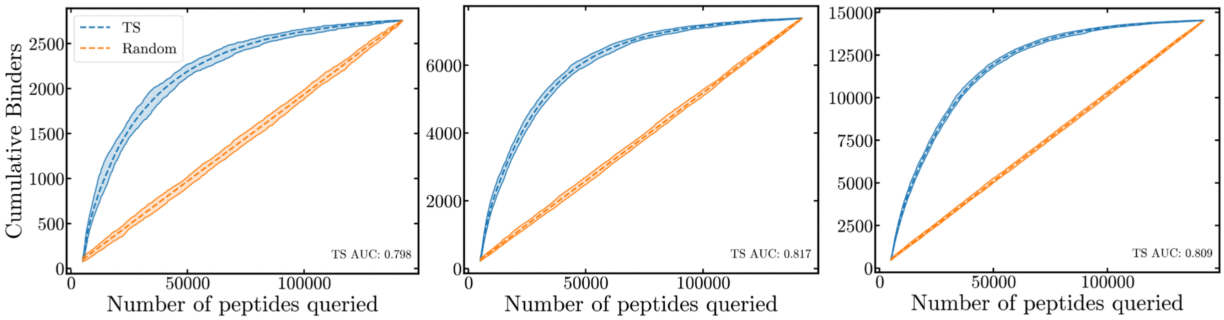


Figure S1: Performance of TS in identifying soluble peptides. Results are shown for three datasets defined by solubility prediction thresholds: peptides with predicted solubility $>$ 0.865 ($\sim$2% of the whole dataset, left), $>$ 0.82 ($\sim$5%, middle), and $>$ 0.77 ($\sim$10%, right) are considered soluble.

### Hyperparameter Screening

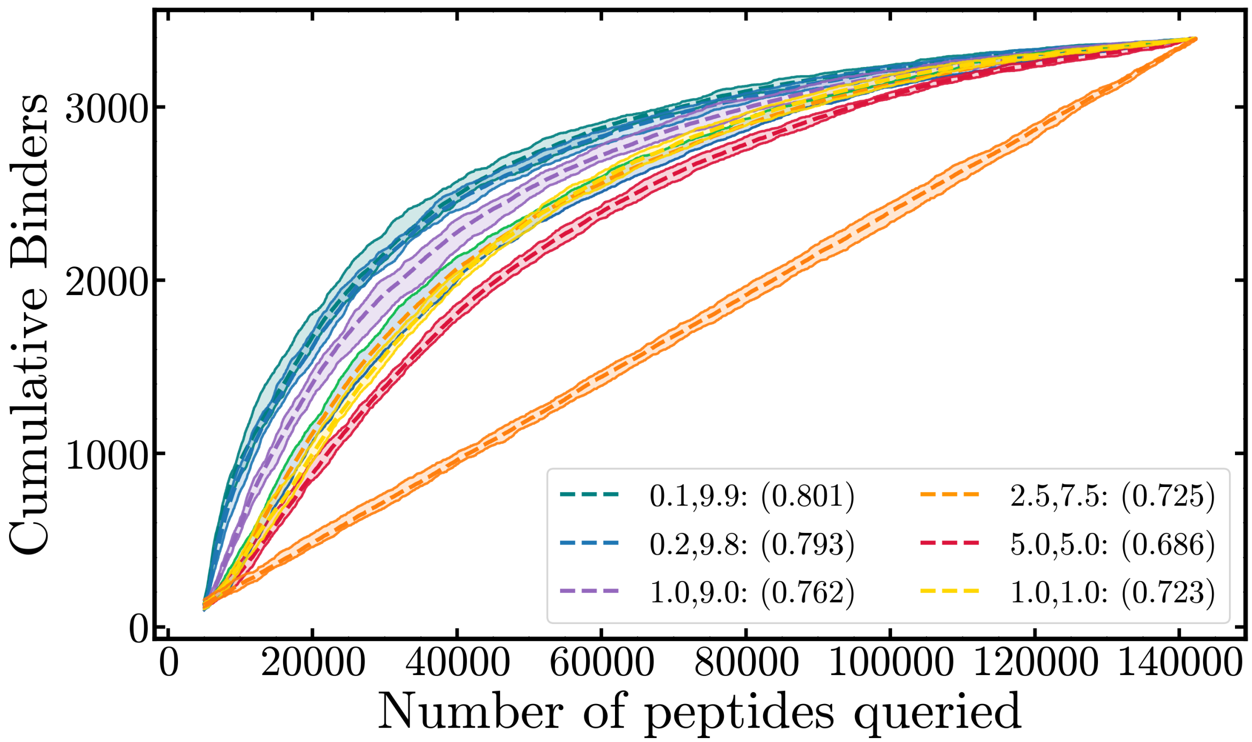


Figure S2: Dependence on the initial priors.

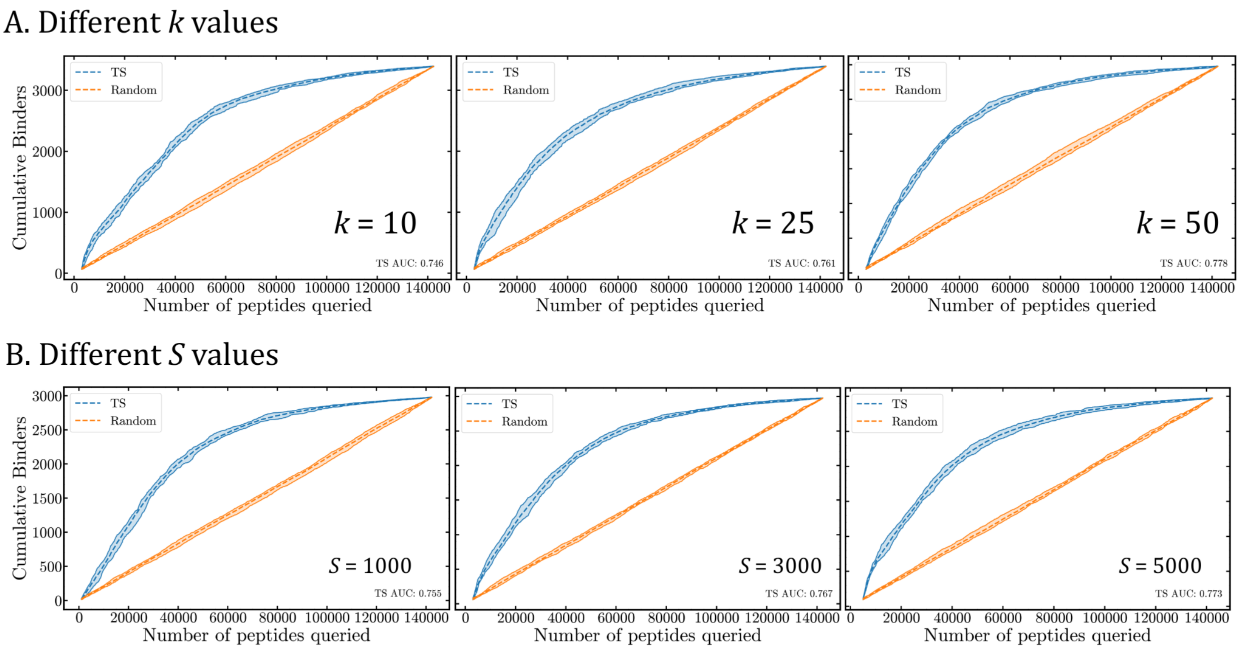


Figure S3: Hyperparameter screening of the (A) numbers of clusters to query and the (B) number of seed sequences with *proportional* allocation.

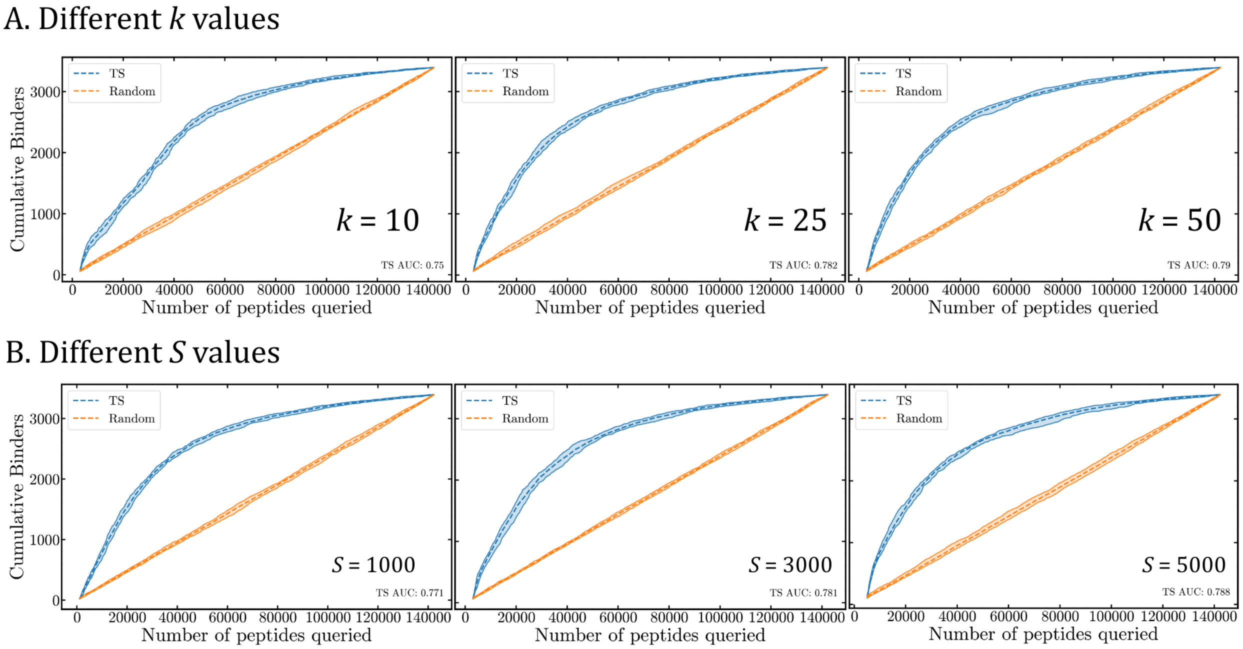


Figure S4: Hyperparameter screening of the (A) numbers of clusters to query and the (B) number of seed sequences with *equal* allocation.

### Distribution of Binders

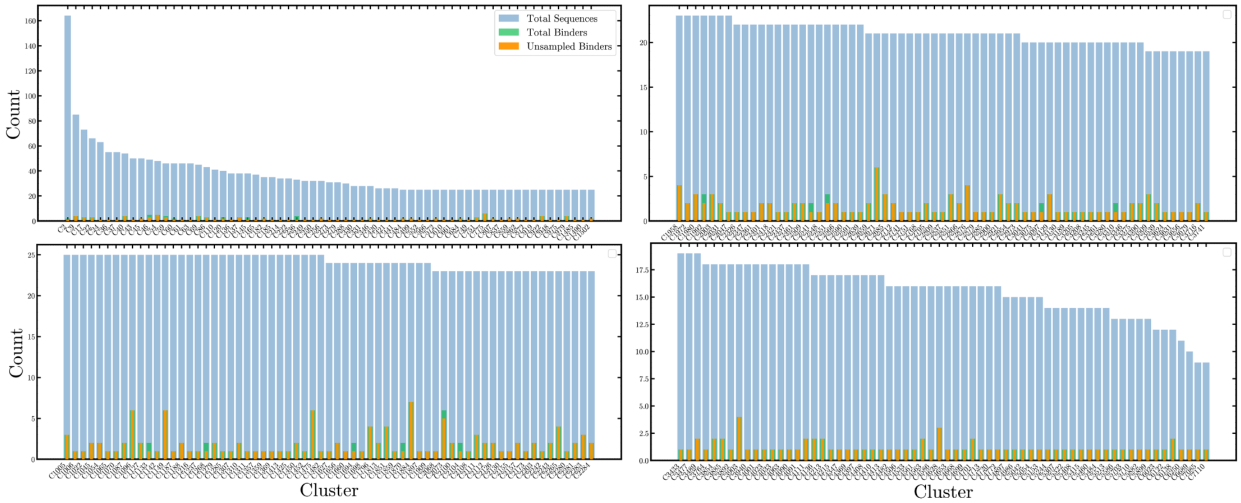


Figure S5: Unsampled clusters after 70k queries

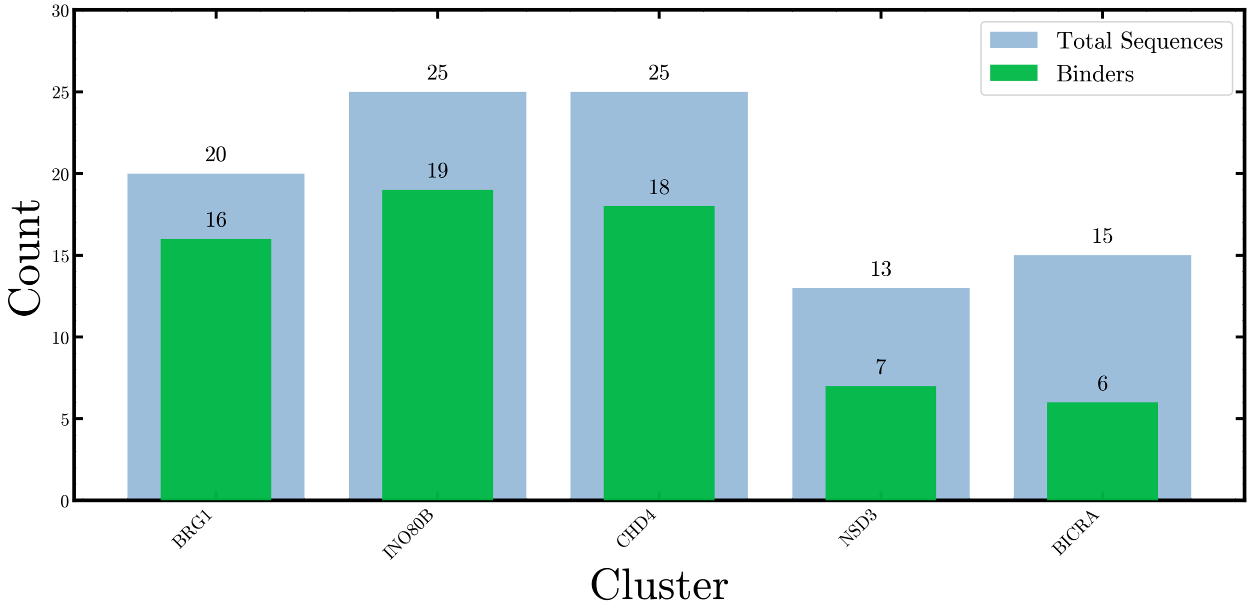


Figure S6: Clusters that contain the top peptides using the CD-HIT algorithm.

### Top Peptides

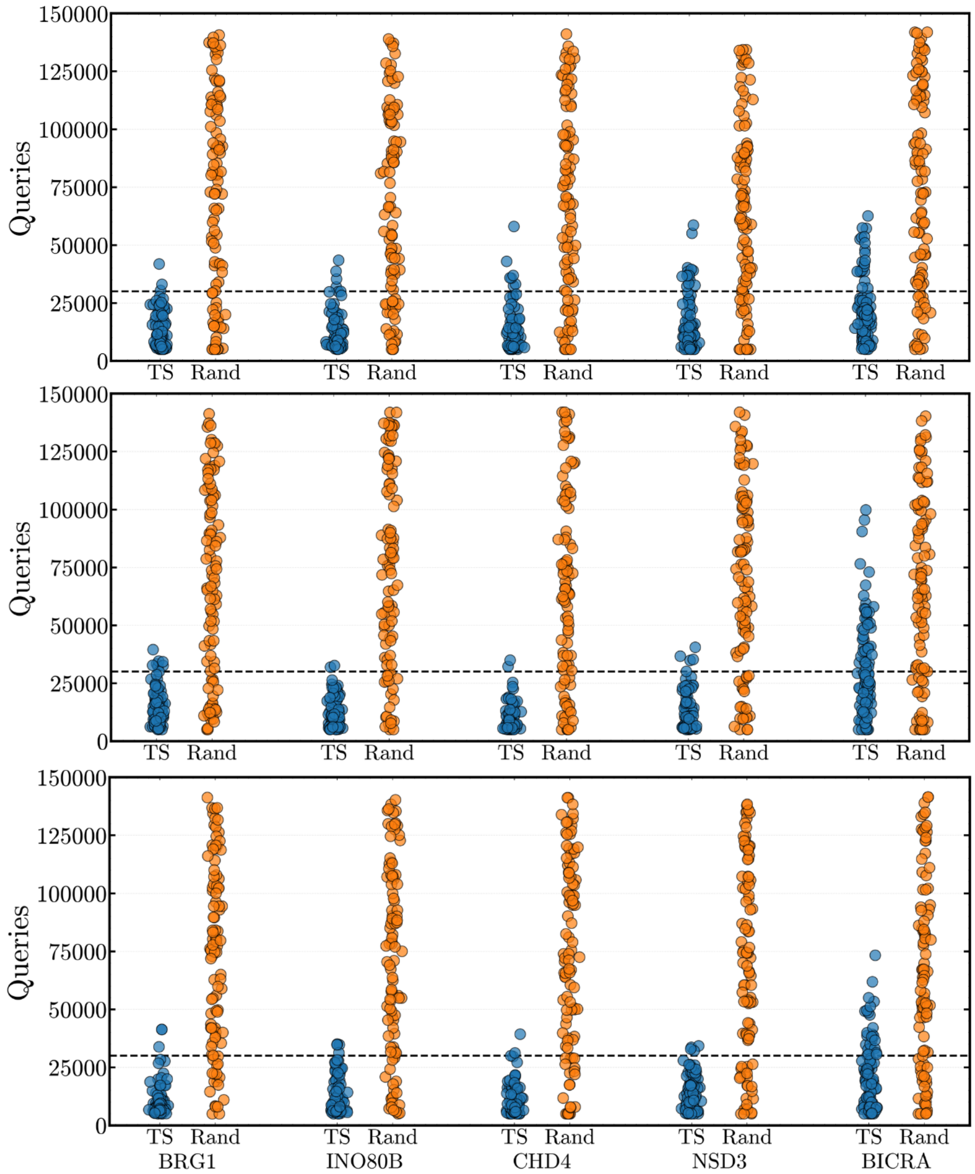


Figure S7: Performance of TS in identifying top peptides using the (top) CD-HIT, (B) Easy-cluster, and (C) Easy-linclust clustering methods. For all results, we used the clusters obtained with sequence identity parameter of 0.5.
